# Requirement and functional redundancy of two large ribonucleotide reductase subunit genes for cell cycle, chloroplast biogenesis in tomato

**DOI:** 10.1101/2020.05.18.102301

**Authors:** Mengjun Gu, Yi Liu, Man Cui, Huilan Wu, Hong-Qing Ling

**Author notes:** Corresponding authors E-mail address (Hong-Qing Ling) E-mail address (Huilan Wu). The author responsible for the distribution of materials in integral to the findings presented in this article in accordance with the policy described in the Instructions for Authors (www.plantcell.org) is: Hong-Qing Ling.

## Abstract

Ribonucleotide reductase (RNR), functioning in the *de novo* synthesis of dNTPs, is crucial for DNA replication and cell cycle progression. However, the knowledge about the RNR in plants is still limited. In this study, we isolated *ylc1* (*<u>y</u>oung <u>l</u>eaf <u>c</u>hlorosis <u>1</u>*) mutant, which exhibited many development defects such as dwarf stature, chlorotic young leaf, and smaller fruits. Map-based cloning, complementation, and knocking-out experiments confirmed that *YLC1* encodes a large subunit of RNR (SlRNRL1), an enzyme involved in the *de novo* biosynthesis of dNTPs. Physiological and transcriptomic analyses indicate that SlRNRL1 plays a crucial role in the regulation of cell cycle, chloroplast biogenesis, and photosynthesis in tomato. In addition, we knocked out *SlRNRL2* (a *SlRNRL1* homolog) using CRISPR-Cas9 technology in the tomato genome, and found that SlRNRL2, possessing a redundant function with SlRNRL1, played a weak role in the formation of RNR complex due to its low expression intensity. Genetic analysis reveals that SlRNRL1 and SlRNRL2 are essential for tomato growth and development as the double mutant *slrnrl1slrnrl2* is lethal. This also implies that the *de novo* synthesis of dNTPs is required for seed development in tomato. Overall, our results provide a new insight for understanding the SlRNRL1 and SlRNRL2 functions and the mechanism of *de novo* biosynthesis of dNTPs in plants.

## Introduction

The balanced supply of the four dNTPs (dATP, dCTP, dGTP, and dTTP), which are building-blocks of DNA synthesis, is critical to the fidelity of faithful genome duplication (Poli et al., 2012). dNTPs can be generated by *de novo* synthesis and salvage pathways. In the salvage pathway, deoxyribonucleosides (dNs) are converted into the corresponding 5′-monophosphate dNs by deoxyribonucleoside kinases (dNKs), and then form dNTPs after phosphorylation. In the pathway of *de novo* biosynthesis, dNTPs are synthesized from simple molecules (Reichard, 1988). The reduction of the four ribonucleoside diphosphates (NDPs) to their corresponding dNDPs catalyzed by ribonucleotide reductase (RNR) is a rate-limiting step in the *de novo* synthesis pathway of dNTPs (Stasolla et al., 2003; Mathews, 2014). The dATP, dCTP, dGTP are directly formed after the phosphorylation of dADP, dCDP, and dGDP by nucleoside diphosphate kinase (NDPK) (Reichard, 1988; Eriksson et al., 2002). Besides of RNR and NDPK, deoxycytidine monophosphate (dCMP) deaminase, thymidylate synthase, and thymidylate phosphate kinase (TMPK) are required for dTTP biosynthesis (Reichard, 1988; Mathews, 2006; Aye et al., 2015).

Eukaryotic RNR is a tetramer composed of two large subunits (R1) and two small subunits (R2) (Elledge et al., 1993). The large subunit of RNR harbors two allosteric sites (activity and specificity sites). Binding ATP to the activity site activates RNR, while binding dATP to the activity site inhibits the activity of RNR (Brown and Reichard, 1969). The activity site regulates the size of the dNTP pool by monitoring the dATP/ATP ratio (Reichard, 1988). The allosteric specificity site is responsible for the balance of the four dNTPs (Brown and Reichard, 1969). The small subunit R2 contains a characteristic diferric-tyrosyl radical cofactor, which is necessary for enzyme activity (Fontecave, 1998). During the reduction reaction, the tyrosyl radical of R2 is transferred to the active site of R1 (Stubbe and Riggs-Gelasco, 1998).

A high concentration of dNTPs is needed in the S-phase of the cell cycle where the DNA duplication takes place, while the requirement of dNTPs is very low in the other phases of the cell cycle (Chabes et al., 2003). Deficiency or imbalance of dNTPs will impede DNA synthesis and lead to a reduced rate of cell proliferation. The RNR activity varies clearly in co-ordination with the demand of dNTPs. Among the cell cycle phases, the activity of RNR is highest in the S phase and lowest in the G_0_ phase (Lowdon and Vitols, 1973). In budding yeast, four genes are involved in RNR formation. *RNR1* and *RNR3* encode R1 protein, while *RNR2* and *RNR4* encode R2 protein (Elledge and Davis, 1987, 1990; Wang et al., 1997; Domkin et al., 2002). The mammalian RNR is encoded by three genes. *RRM1* encodes the large subunit, whereas *RRM2* and *RRM2B* encode distinct small subunit R2 and p53R2, respectively (Guittet et al., 2001; Xue et al., 2003; Qiu et al., 2006). Previous studies showed that the *RNR1* mRNA level fluctuated more than tenfold and the *RNR2* mRNA level varied only about twofold during the cell cycle of yeast, the most abundance of *RNR1* and *RNR2* mRNA was observed in the S phase of the cell cycle (Elledge et al., 1992). In mammalian cells, RRM1 expressed constantly, while the RRM2 expression was specifically in S phase, and p53R2 expression at a low level was constant in all stages of its cell cycle (Björklund et al., 1990).

R1 and R2 are necessary for cells to enter mitosis in yeast because loss of their function in mutant *rnr1* and *rnr2* results in lethality (Elledge and Davis, 1987, 1990). Deletion of *RNR4*, which encodes a R2 protein, resulted in the imbalance of the dNTP pools and a slow growth rate at optimal growth temperature, whereas the cell division of the mutant was arrested in the S phase of the cell cycle at the restrictive temperature (Huang and Elledge, 1997; Wang et al., 1997). It was also demonstrated that inhibition of the RNR activity with hydroxyurea slowed down the DNA replication of budding yeast and arrested its cell cycle in the S phase (Barlow et al., 1983; Alvino et al., 2007). Wanrooij and Chabes reported that RNR was crucial not only for nuclear DNA replication, but also for the replication of mitochondrial DNA (mtDNA) (Wanrooij and Chabes, 2019). In the yeast, *RNR4 is* required for the maintenance of the mitochondrial genome because its mutant *rnr4* lacked functional mitochondria. In human and mouse, the mutation of *RRM2B* led to mtDNA depletion and resulted in mitochondrial disease (Powell et al., 2005; Bourdon et al., 2007; Bornstein et al., 2008; Chimploy et al., 2013; Xue et al., 2015).

In plants, mutations of *RNR* genes result in dwarf plants, abnormal leaf development and defective chloroplasts (Wang and Liu, 2006; Garton et al., 2007; Yoo et al., 2009; Qin et al., 2017; Tang et al., 2019). *Arabidopsis thaliana cls8* and *tso2* mutants, defective in the large subunit and the small subunit of RNR, respectively, exhibited bleached leaves and siliques (Wang and Liu, 2006; Garton et al., 2007). Additionally, the chloroplast number of *cls8* reduced, and the chloroplasts were dysplastic (Garton et al., 2007). In contrast to Arabidopsis, the rice mutant *v3* with a disrupted large subunit gene and *st1* with a disrupted small subunit gene showed growth-stage-specific and environment-dependent leaf chlorosis (Yoo et al., 2009). Similarly, the *Setaria italica* mutant *sistl1* showed growth retardation, striped leaf phenotype and reduction of its chloroplast biogenesis due to the lost function of the large subunit of RNR (Tang et al., 2019). Although several studies have been done in plants, the knowledge of RNR functions is still limited.

In this study, null mutants of *SlRNRL1* and *SlRNRL2*, which encode the large subunit of the RNR complex in tomato, have been identified and generated. With them, we studied the biological functions of SlRNRL1 and SlRNRL2, and demonstrated that SlRNRL1 was more important than SlRNRL2 in the *de novo* biosynthesis of dNTPs in tomato. Furthermore, we also showed that *SlRNRL1* and *SlRNRL2* are required for tomato growth and development.

## Results

### Phenotypic characterization of *ylc1* mutant

The *ylc1* (*<u>y</u>oung <u>l</u>eaf <u>c</u>hlorosis <u>1</u>*) mutant was identified from the mutation population of the tomato cultivar Moneymaker (MM) mediated by ethyl methyl sulfonate (EMS). Its young leaves exhibited chlorosis (Figure 1A). Along with the growth, the chlorotic leaves became green (Figure 1B). In accordance with the chlorotic phenotype, the content of total chlorophyll, chlorophyll a, and chlorophyll b of *ylc1* was significantly lower than that of MM (Figure 1C, 1D). Compared to MM, the *ylc1* mutant displayed smaller leaves and shorter plant height (Figure 1B, 1E), and the fresh and dry weights of its roots and shoots were markedly declined (Figure 1F, 1G). In addition to the leaf color and biomass, the fruit of the *ylc1* mutant was also significantly smaller than that of MM (Figure 1H, 1I). These results indicated that *YLC1* plays some important roles in tomato growth and development.

**Figure 1.**
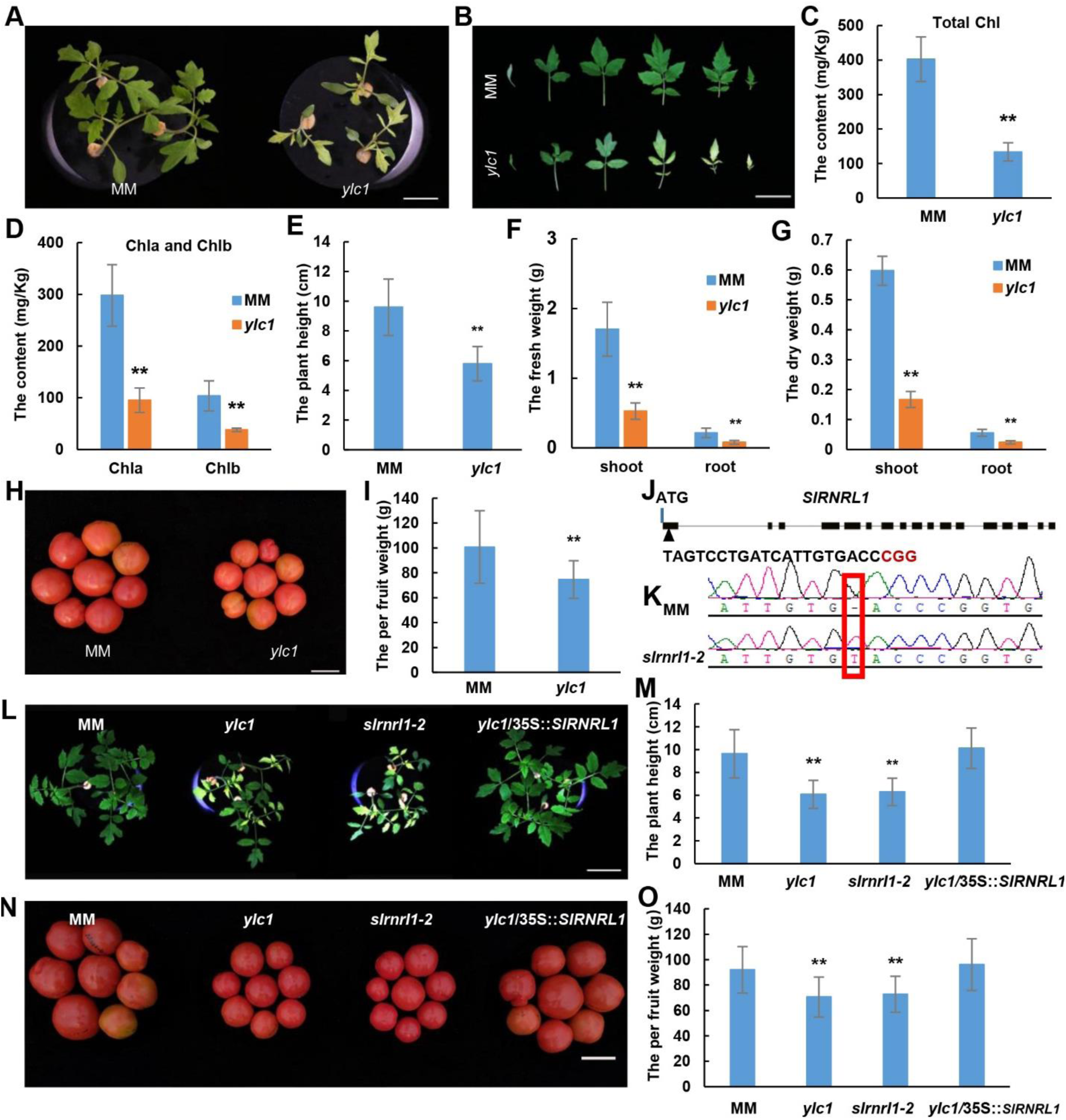
Phenotypic characterization of *ylc1* and the functional verification of *YLC1*. (**A**) The phenotype of MM and *ylc1* mutant grown in the glasshouse for 10 days. (B) Leaves of MM and *ylc1* plants grown in the glasshouse for 21 days. (C-D) Content of total chlorophyll, chlorophyll a and chlorophyll b of young leaves. (E) The plant height of MM and *ylc1* plants grown in the glasshouse for 21 days. (F), The plant fresh weight of MM and *ylc1* grown in the glasshouse for 21 days. (G) The dry weight of five plants grown in the glasshouse for 21 days. (H-I) The fruit phenotype of MM and *ylc1*. (J) The gene structure of *SlRNRL1* and the sgRNA (single guide RNA) sequence of the CRISPR construct. The black boxes stand for the exons and black lines represent introns. The PAM (protospacer adjacent motif) sequence is highlighted in red. (K) The sequence of the target regions in *SlRNRL1* of MM and the nucleotide insertion marked with red box. (L-M) The plant phenotype of the MM, *ylc1*, the knocking-out mutant *slrnr1-2*, and complemented plants (*ylc1*/35S::*SlRNRL1*). (N-O) The fruit phenotype of MM, *ylc1*, *slrnr1-2* and complemented plants. The means and standard deviations were obtained from three biological replicates. ** indicates significant differences by t-test (P < 0.01) compared to MM. Bars, 5 cm.

### *YLC1* encodes the large subunit of RNR complex

For genetic mapping of the *ylc1* mutant, we crossed *ylc1* with a wild species, *Solanum pennellii* (LA0716). The F1 plants grew normally with green leaves. In F2 generation, the segregation of the mutant and wild type appeared at a ratio of 1:3. These demonstrated that the mutant phenotype of *ylc1* was caused by a recessive mutation at a single nuclear gene.

Through bulk segregation analysis and whole-genome resequencing of F2, the *ylc1* locus was delimited to a 1.5 Mb region from 3.5 Mb to 5.0 Mb on chromosome 4 (Supplemental Figure 1A). To finely map this region, INDEL markers were developed by comparing the genomic sequences of LA0716 and *ylc1*. Using the INDEL markers, the *YLC1* gene has been narrowed to an 86.76 kb region from 4,380,741 bp to 4,467,501 bp between INDEL-7 (M7) and INDEL-8 (M8) on chromosome 4 (Supplemental Figure 1B). In this region, there are 12 genes predicted. Among them, only one mutation site with 2-bp deletion in the fourteenth exon of *Solyc04g012060* has been identified by gene amplification and sequencing (Supplemental Figure 1C). The 2-bp deletion led to a frame shift and premature termination of the predicted protein. According to the tomato genome annotation (https://www.solgenomics.net/), *Solyc04g012060* encodes a large subunit of RNR. Sequence analysis showed that another gene, *Solyc04g051350*, which shows 92.95% identity with *Solyc04g012060*, is present in the tomato genome. Therefore, we termed *Solyc04g012060* as *SlRNRL1* (*<u>r</u>ibo<u>n</u>ucleotide <u>r</u>eductase <u>l</u>arge subunit <u>1</u>*) and *Solyc04g051350* as *SlRNRL2* (*<u>r</u>ibo<u>n</u>ucleotide <u>r</u>eductase <u>l</u>arge subunit <u>2</u>*).

To confirm whether *SlRNRL1* is the target gene, we constructed a CRISPR-CAS9 vector targeting exon 1 of *SlRNRL1* (Figure 1J), introduced it in the genome of MM by Agrobacterium-mediated transformation, and obtained a homozygous transgenic line (termed as *slrnrl1-2*), in which the *SlRNRL1* was knocked out by insertion of a T nucleotide at the 104^th^ bp position of the coding region of *SlRNRL1* (Figure 1K). Like the *ylc1* mutant, *slrnrl1-2* exhibited chlorosis on its young leaves and retarded growth (Figure 1L, 1M). Similar to *ylc1*, its fruit size was smaller than that of the wild type (Figure 1N, 1O). In parallel, we performed a genetic complementation experiment. The transformation vector with a 2727-bp length of *SlRNRL1* CDS under the control of CaMV35S promoter was generated and introduced into the *ylc1* mutant by *Agrobacterium tumefaciens*. The mutant phenotype was completely rescued in the transgenic lines expressing *SlRNRL1* (Figure 1L-1O). Taken all together, the defect of *SlRNRL1* is responsible for the *ylc1* phenotype. Thus, we renamed the mutant *ylc1* as *slrnrl1-1* here after.

### The expression pattern of *SlRNRL1* and subcellular localization of its encoded protein

To characterize the expression profiles of *SlRNRL1*, we did quantitative reverse transcription-PCR (qRT-PCR) analysis. As shown in Figure 2A, *SlRNRL1* expressed in tissues including root, stem, leaf, and flower. The highest expression intensity was detected in the leaf among the tissues examined. Furthermore, we generated transgenic tomato plants harboring a *GUS* reporter gene driven by the *SlRNRL1* promoter (p*SlRNRL1*::*GUS*). Histochemical assays showed that the *GUS*, consistent with the expression profiles of *SlRNRL1*, expressed in the tips of primary root and lateral root, lateral root, primordium, leaf, and flower (Figure 2B).

**Figure 2.**
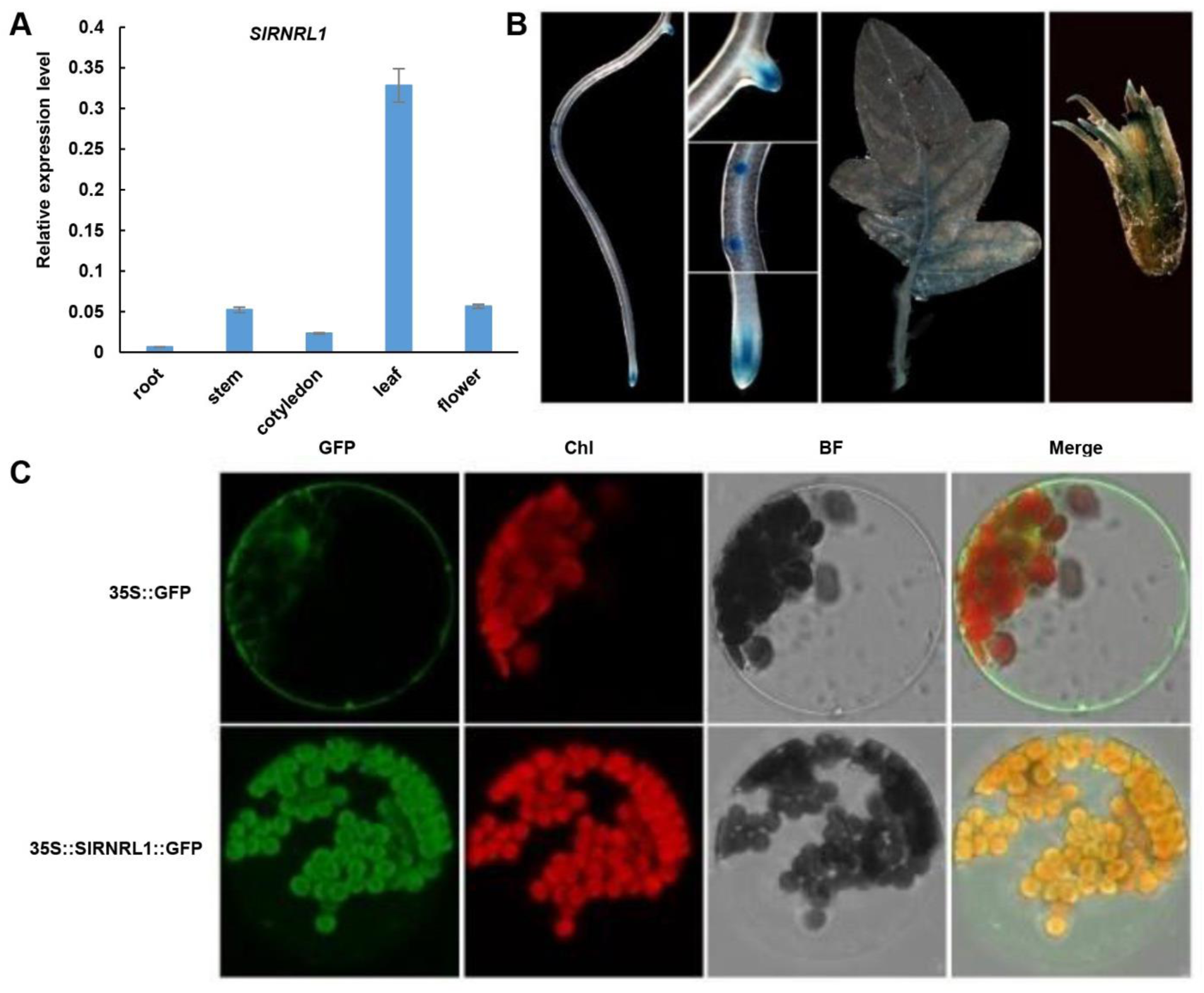
The expression profiles of *SlRNRL1* and subcelluar localization of SlRNRL1. (A) The relative expression level of *SlRNRL1* in the root, stem, cotyledon, leaf, and flower. *Kinase I* was used as the internal control. The means and standard deviations were obtained from three independent replicates. (B) Histochemical assay of *SlRNRL1* expression patterns. (C) Subcellular localization of SlRNRL1 in Arabidopsis protoplast. GFP, green fluorescent protein; Chl, chlorophyll; BF, bright field.

To examine the subcellular localization of SlRNRL1, the expression vector containing a *SlRNRL1-GFP* fusion protein gene controlled by the CaMV35S promoter was constructed and transiently expressed in the protoplasts of Arabidopsis. As shown in Figure 2C, the GFP signal was clearly co-localized with the signal of chlorophyll. These results indicated that SlRNRL1 should be localized in the chloroplasts of leaves or in plastids of other tissues.

Further, we analyzed the protein structure of SlRNRL1 using the Interpro protein prediction database (http://www.ebi.ac.uk/interpro/). As shown in Supplemental Figure 2A, SlRNRL1 contains three conserved domains, ATP cone domain (amino acids 1–90), R1 N-terminal domain (amino acids 142–212), and R1 C-terminal domain (amino acids 757–757). Alignment of amino acid sequences revealed that SlRNRL1 is conserved and exhibits a high similarity to R1 of other plant species (Supplemental Figure 2B).

### The loss function of *SlRNRL1* affects cell cycle progress, chloroplast division, and photosynthesis capacity

As RNR functions in the biosynthesis of dNTPs, which are required for DNA duplication, the cell division of *slrnrl1* mutant may be aberrant. To check the hypothesis, we performed histological transparency analysis and found that the mesophyll cell number of *slrnrl1-1* was significantly less than that of MM (Figure 3A; Supplemental Figure 3A, 3B). Flow cytometric analysis revealed that the percentage of 4C and 8C cells in *slrnrl1-1* was markedly lower than that of MM (Figure 3B; Supplemental Figure 3C, 3D). These results demonstrated that the DNA duplication and progress of cell cycle in *slrnrl1-1* are indeed affected.

**Figure 3.**
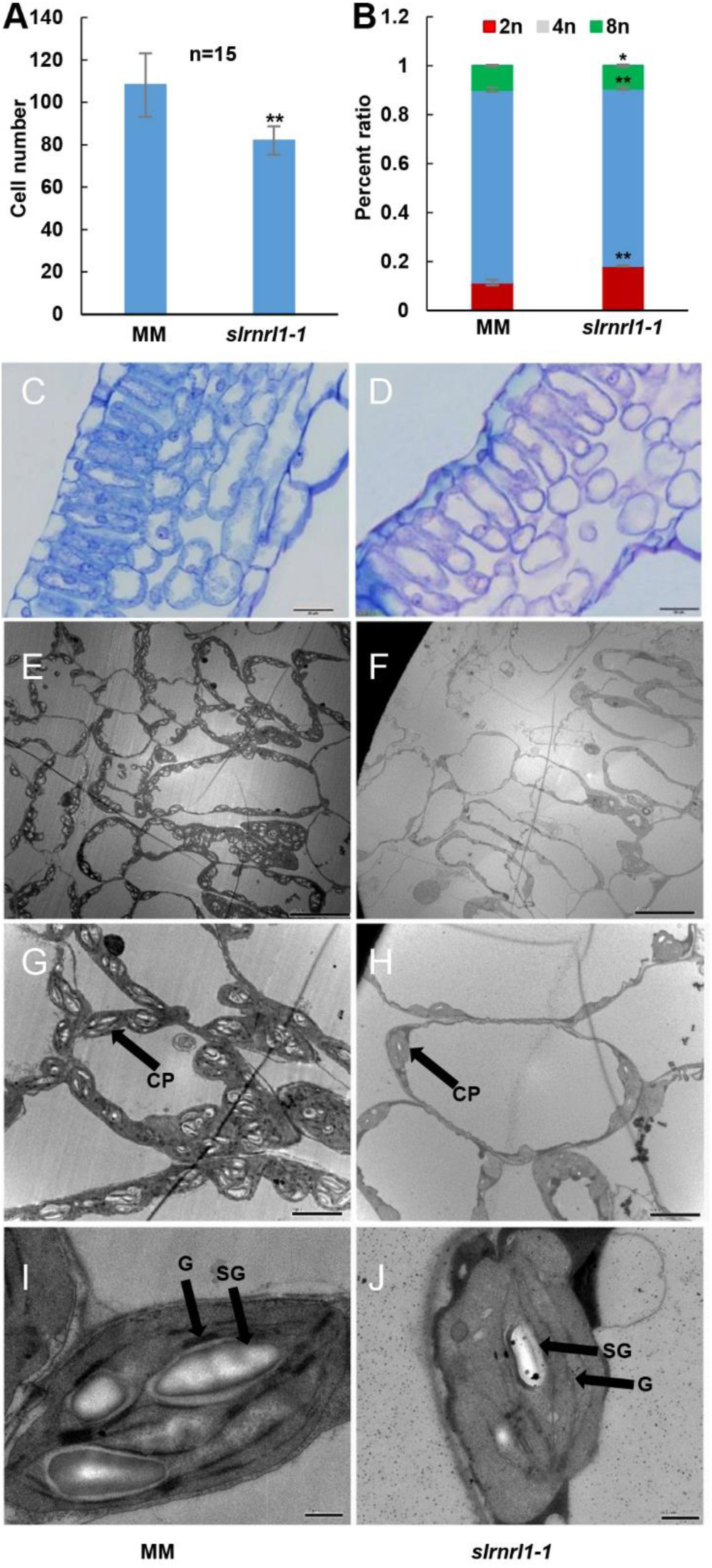
The nuclear ploidy and the microscopy of leaf tissues of MM and *slrnrl1-1*. (A) The number of palisade cells of 6-day-old leaf in MM and *slrnrl1-1*. ** indicates significant differences by t-test (P<0.01) between MM and *slrnrl1-1*. (B). Quantification of the DNA profiles of MM and *slrnrl1-1*. The means and standard deviations were obtained from three biological replicates. ** indicates significant differences by t-test (P < 0.01) between MM and *slrnrl1-1*. * indicates significant differences by t-test (P < 0.05) between MM and *slrnrl1-1*. (C, D) Transverse sections of 6-day-old leaf blades from MM (C) and *slrnrl1-1* (D). Bars, 20 µm. (E-J) Ultrastructure of 6-day-old leaf blades from MM (E, G, I) and *slrnrl1-1* (F, H, J). CP, chloroplast; G, grana; SG, starch granule. Bars, (E, F) 20 µm; (G, H) 5 µm; (J, K) 0.5 µm.

Considering that SlRNRL1 is localized in chloroplasts, we observed the chloroplast number and its ultrastructure via transmission electron microscope, and showed that the number of chloroplasts in *slrnrl1-1* was surely reduced compared to MM (Figure 3C-H). Furthermore, we performed qRT-PCR analysis to detect the ratio of chloroplast DNA to nuclear DNA in MM and *slrnrl1-1*. The result revealed that the ratio in the young leaves of *slrnrl1-1* reduced significantly compared to that of MM (Supplemental Figure 4A). All these data indicate that plastid DNA duplication during chloroplast biogenesis is inhibited in the young leaves of *slrnrl1-1*. Additionally, we observed the chloroplast size using a transmission electron microscope. As shown in Figure 3I and 3J, the size of chloroplasts in *slrnrl1-1* was much smaller than that of MM. The chloroplast thylakoids of *slrnrl1-1* displayed fewer grana stacks than that of MM, although the chloroplasts contained differentiated thylakoids (Supplemental Figure 3I, 3J). These results indicate that deficiency of SlRNRL1 also disrupts chloroplast division and development.

For the investigation of photosynthesis affection, we investigated the number and size of starch grains of leaves. As shown in Figures 3I and 3J, the number and size of starch grains in the leaves of *slrnrl1-1* decreased notably compared to that of MM. I_2–_KI staining revealed that the staining color of MM leaf was deeper than that of *slrnrl1-1* (Supplemental Figure 4B), indicating that less starch was accumulated in *slrnrl1-1* leaves than that of MM. Moreover, we measured the net photosynthetic rate. As shown in Supplemental Figure 4C, the net photosynthetic rate in the leaf of *slrnr1-1* was significantly decreased, compared to MM. All the results imply that photosynthesis capacity is reduced in *slrnrl1-1*.

### Comparison of genome-wide transcriptomes of MM and *slrnrl1-1*

To elucidate the functions of SlRNRL1 in tomato growth and development, we compared the genome-wide transcriptomes of MM and *slrnrl1-1* leaves, and found 219 down-regulated and 1,464 up-regulated genes in the first true leaf of *slrnrl1-1* at 6-days-old stage, compared to MM (Supplemental Figure 5A, S5B). The Gene Ontology (GO) analysis of the 219 down-regulated genes showed that they enriched in GO terms related to pigment binding, photosystem, photosynthesis, and chloroplast development (Figure 4A). The GO analysis of the 1,464 up-regulated genes showed that they enriched in GO terms related to DNA repair, DNA metabolic process, organelle fission, and cell cycle (Figure 4B).

**Figure 4.**
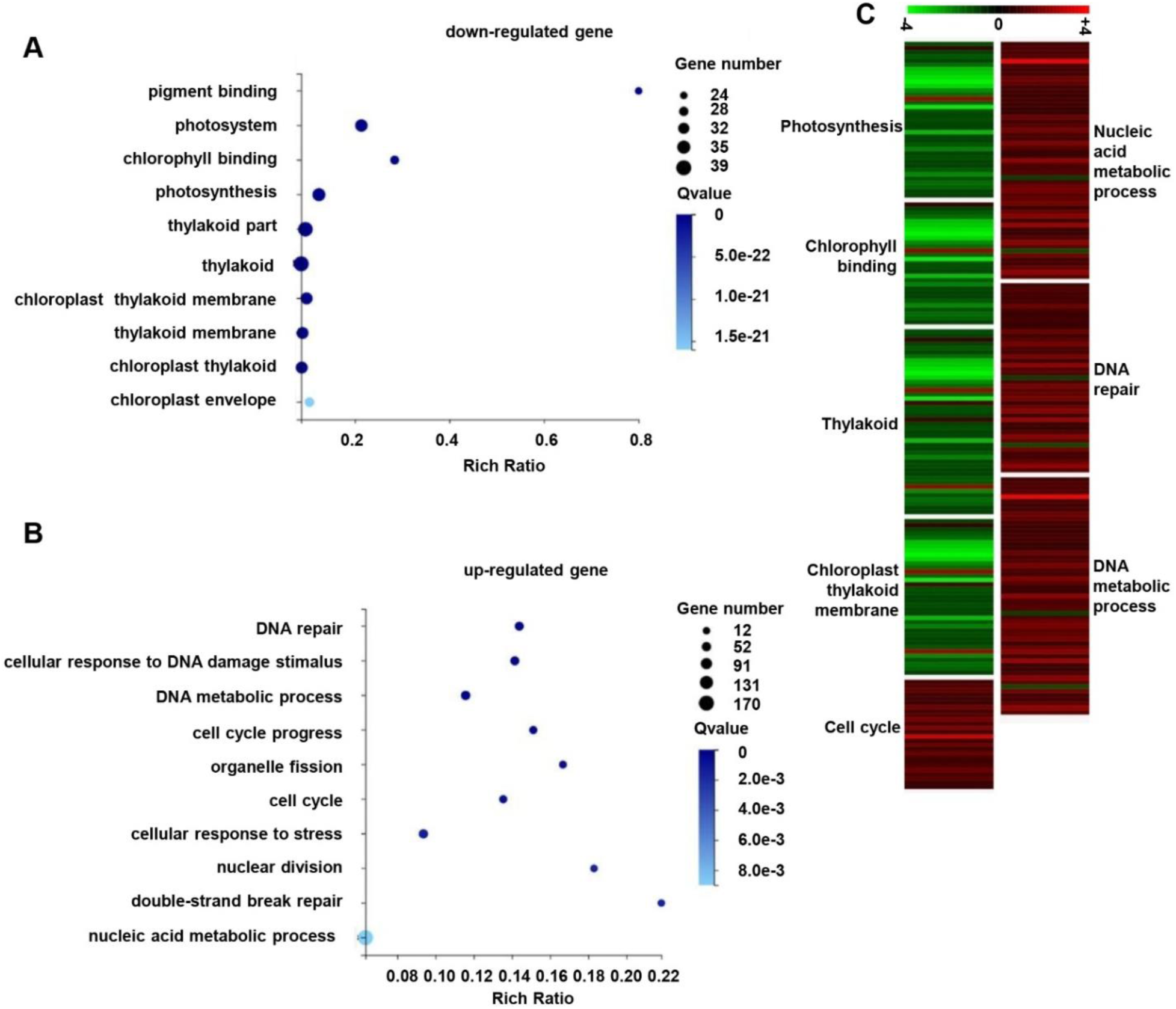
RNA-seq analysis of MM and *slrnrl1-1*. (A, B) GO terms enriched analysis of down-regulated genes (A) and up-regulated genes (B). (C) Heatmaps of DEGs in GO terms associated with mutant phenotype regulation. Each group of color-block represents a GO term. The names of GO terms are shown beside each block group. Color of each block represents the value of log2 (FPKM-*slrnrl1-1*/FPKM-MM). The color bar, from –4 to 4. Data were obtained from three biological replicates.

Given the functions of SlRNRL1 and the phenotypes of *slrnrl1-1*, we selected the GO terms related to photosynthesis, chlorophyll binding, thylakoid, chloroplast thylakoid membrane, nucleic acid metabolism process, DNA repair, DNA metabolic process and cell cycle for deep analysis. Most of the different expression genes (DEGs) related to chloroplast, thylakoid membrane, thylakoid, photosynthesis, and chlorophyll binding were down-regulated (Figure 4C). This was consistent with the abnormal stacking of thylakoids, decreased chlorophyll, reduced number of chloroplasts, and starch grains in *slrnrl1-1* (Figure 3C-J, 1C, 1D; Supplemental Figure 4B). Almost all the DEGs in nucleic acid metabolism process, DNA repair pathway, DNA metabolic process, and cell cycle were up-regulated (Figure 4C). This may be caused by feedback regulation of the functional defect of SlRNRL1. For validation of the RNA-seq results, we selected 6 genes, which are associated with photosynthesis (*Solyc02g071010*), chlorophyll binding (*Solyc03g115900*), thylakoid (*Solyc07g063600*), cell cycle (*Solyc10g074720*), DNA repair pathway (*Solyc05g053520*) and DNA metabolic process (*Solyc03g117960*), and checked their expression level by qRT-PCR analysis. The expression abundance of the six genes was consistent with that detected in RNA-seq (Supplemental Figure 6A, 6B).

### SlRNRL2 is a SlRNRL1 homolog with a weak function

Phylogenetic analysis showed that R1 proteins with a close relationship are widely present in dicot and monocot plants (Figure 5A). *SlRNRL1* and *SlRNRL2* are two genes encoding R1 protein in the tomato genome, sharing a high sequence identity (92.95%) each other at the protein level. The two genes possess the same gene structure containing 17 exons and 16 introns. As shown in Figure 5B, *SlRNRL2* revealed a similar expression profile as *SlRNRL1*, but its expression intensity was much lower.

**Figure 5.**
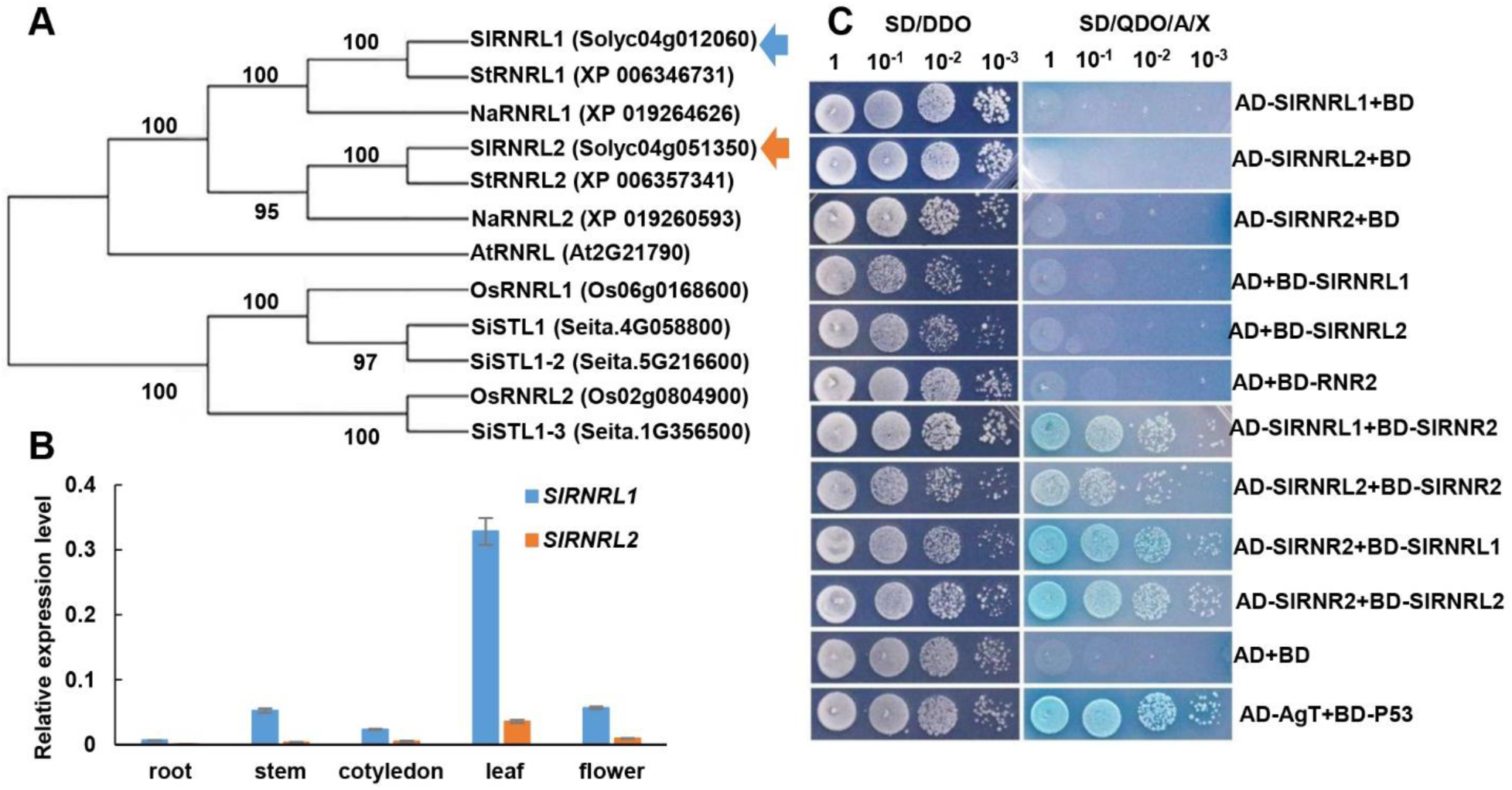
The homologous gene of *SlRNRL1* in tomato. (A) Phylogenetic analysis of R1 proteins in eudicot tomato (*S. lycopersicum*), potato (*S. tuberosum*), tobacco (*Nicotiana attenuata*), Arabidopsis (*A. thaliana*), and in monocot rice (*Oryza sativa*), millet (*Setaria italica*). The phylogenetic tree was constructed using MEGA 4. Sl, *S. lycopersicum*; St, *S. tuberosum*; Na, *N. attenuate*; At, *A. thaliana*; Os, *O. sativa*; Si, *S. italica*. Blue arrow indicates the SlRNRL1 protein, yellow arrow indicates SlRNRL2. (B) qRT-PCR analysis of *SlRNRL1* and *SlRNRL2* in the root, stem, cotyledon, leaf, and flower. *Kinase I* was used as the internal control. The means and standard deviations were obtained from three biological replicates. (C) Yeast two-hybrid analysis of SlRNRL1, SlRNRL2 and SlRNR2. ×10^−1^, yeast diluted 10 times; ×10^−2^, yeast diluted 100 times; ×10^−3^, yeast diluted 1000 times. AD, pGADT7 vector; BD, pGBKT7 vector. DDO, SD-Leu-Trp; QDO/X/A, SD-Leu-Trp-His + Ade + X-α-Gal + Aureovasidin A.

To verify whether the proteins encoded by *SlRNRL1* and *SlRNRL2* are involved in the formation of RNR complex, the full-length coding sequences of *SlRNRL1* and *SlRNRL2* (encoding the large subunit of the RNR complex), and *SlRNR2* (encoding a small subunit of the RNR complex) were cloned into pGBKT7 and pGADT7, and expressed in yeast. The yeast cells co-transformed with *SlRNR2*-BD and *SlRNRL1*-AD, *SlRNR2*-BD and *SlRNRL2*-AD, *SlRNR2*-AD and *SlRNRL1*-BD, *SlRNR2*-AD and *SlRNRL2*-BD grew well on both double dropout medium (DDO) and quadruple dropout medium (QDO) supplemented with X-α-Gal and aureobasidin A (Figure. 5C). These results demonstrate that both SlRNRL1 and SlRNRL2 are able to interact with SlRNR2 and are involved in RNR complex formation.

To characterize the function of SlRNRL2, we then generated a loss-of-function mutant of SlRNRL2 (*slrnrl2*) using the CRISPR/Cas9 system, in which a single T-nucleotide was inserted at the position of 1219 bp in the coding region of *SlRNRL2*. The insertion of the T nucleotide led to a frame shift and premature termination of SlRNRL2 protein in *slrnrl2* (Figure 6A, 6B). Unlike *slrnrl1*, the *slrnrl2* mutant did not exhibit any visible phenotype (Figure 6C, 6D). Furthermore, we determined the relative expression level of *SlRNRL2* in the leaf of *slrnrl1-1* mutant at different development stages. As shown in Figure 7E, a similar expression level of *SlRNRL2* was detected in the leaves of MM and *slrnrl1-1* at the early development stage (6-days old), whereas *SlRNRL2* showed notably higher expression in leaves of *slrnrl1-1* than that of MM at the later stages (from 8-day-old to 14-day-old). The higher expression intensity of *SlRNRL2* in the later stages may be the reason for the leaf color recovery (green) of *slrnrl1-1* along with its growth. These results indicate that SlRNRL2 has redundant functions as SlRNRL1 and plays a weak role in the formation of tomato RNR complex due to its low expression intensity (Figure 5B).

**Figure 6.**
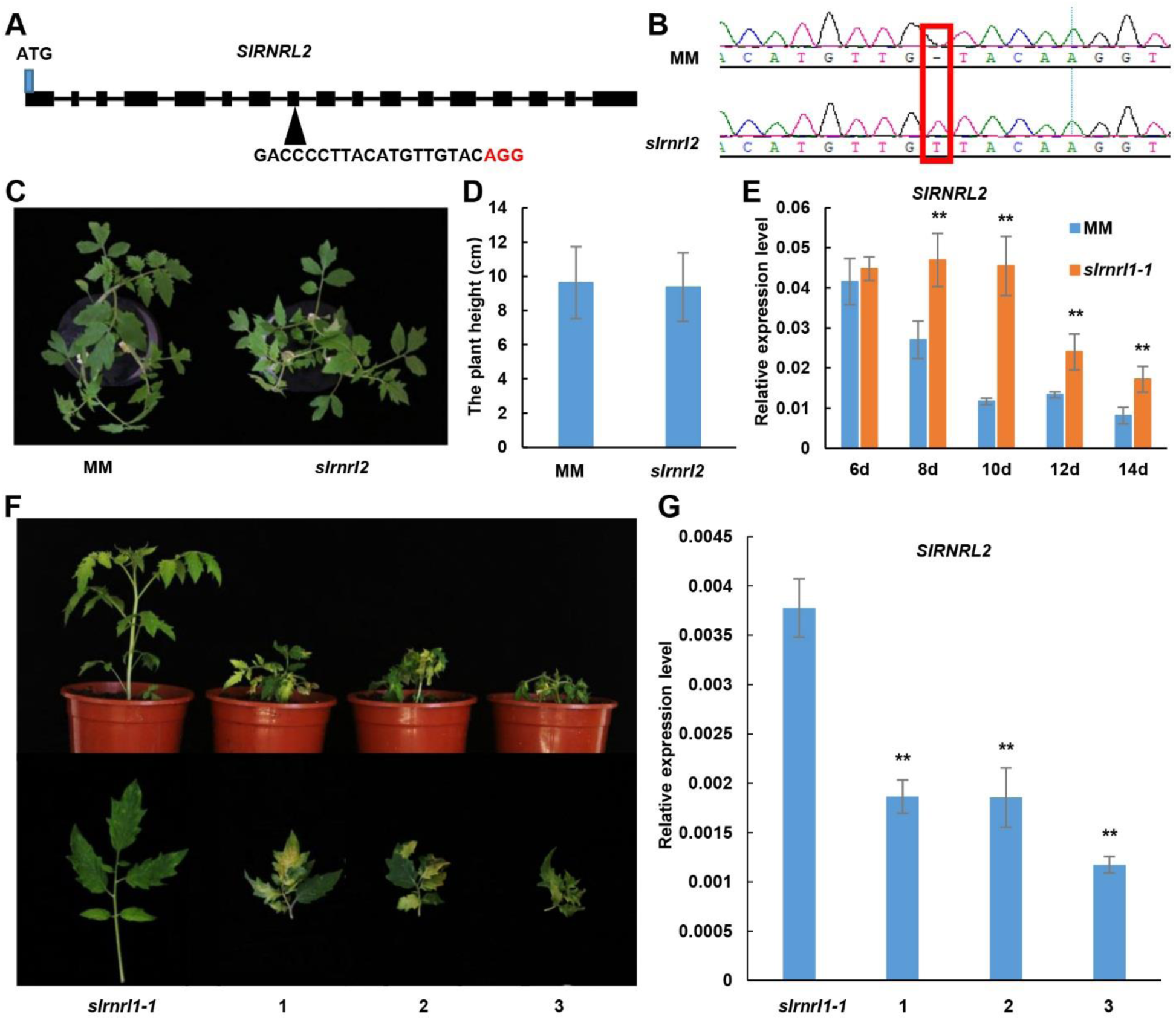
Knocking-out of *SlRNRL2* in MM and Silencing of *SlRNRL2* in *slrnrl1-1*. (A) The gene structure of *SlRNRL2* and the sgRNA sequence of the CRISPR construct. The black boxes stand for the exons and black lines represent introns. The PAM sequence is highlighted in red. (B) The sequences of the target regions in MM and a nucleotide insertion marked with red box. (C) Phenotype of MM and the knocking-out mutant *slrnrl2* grown in the glasshouse for 21 days. (D) The plant height of MM and *slrnrl2* grown in the glasshouse for 21 days. (E) qRT-PCR analysis of *SlRNRL2* expression in leaves of MM and *slrnrl1-1*. 6d, 8d, 10d, 12d, 14d represent the 6-day-old leaf, 8-day-old leaf, 10-day-old leaf, 12-day-old leaf and 14-day-old leaf, respectively. *Kinase I* was used as the internal control. The means and standard deviations were obtained from three biological replicates. (F) The phenotype of *slrnrl1-1* and *slrnrl1-1* infiltrated pTRV1/pTRV2-*SlRNRL2*. (G) qRT-PCR analysis of *SlRNRL2* expression in leaves of *slrnrl1-1* and *SlRNRL2*-silenced *slrnrl1-1* plants (1, 2, 3). Values are the mean of 3 replicates ±s.d. ** indicates significant differences by t-test (P < 0.01) between *slrnrl1-1* and *SlRNRL2*-silenced *slrnr1-1* plants.

### SlRNRL1 and SlRNRL2 are required for tomato growth and development

To further characterize the functions and requirements of SlRNRL1 and SlRNRL2 for tomato growth and development, we crossed *slrnrl1-1* and *slrnrl2* and tried to generate the double mutant *slrnrl1-1rnrl2*. The F1 plants grew normally as wild type. In F2 population, we analyzed 272 individual plants and detected all expected-genotypes except for the double mutant genotype *slrnrl1-1rnrl2* (Table 1). Considering that some seeds of F2 population have not germinated, we then analyzed the un-germinated seeds and still did not identify any seed with the double mutant *slrnrl1-1rnrl2* genotype (Table 1). These results strongly suggest that the double mutation of *SlRNRL1* and *SlRNRL2* should be lethal before or during seed development.

**Table 1.** Genotyping of F2 populations derived from the cross between *slrnrl1-1* and *slrnrl2*. “A” means the allele of *SlRNRL1*, “a” refers to the allele of *slrnrl1-1*; “B” means the allele of *SlRNRL2*, “b” refers to the allele of *slrnrl2*.

| Genotype | Number of plants | Number of<br>none-germinated<br>seeds | Total number |
| --- | --- | --- | --- |
| AABB | 3 | 1 | 4 |
| AABb | 12 | 2 | 14 |
| AAbb | 67 | 8 | 75 |
| AaBB | 3 | 2 | 5 |
| AaBb | 110 | 16 | 126 |
| Aabb | 32 | 1 | 33 |
| aaBB | 12 | 1 | 13 |
| aaBb | 33 | 4 | 37 |
| aabb | 0 | 0 | 0 |
| Total | 272 | 35 | 307 |

Furthermore, we tried to down-regulate *SlRNRL2* expression in the genetic background of *slrnrl1-1* using a tobacco rattle virus-induced gene silencing (VIGS) system. Firstly, we tested the VIGS system by infiltrating the leaves of MM with pTRV1/pTRV2-*SlPDS* (as a positive control). As shown in Supplemental Figure 7A, a photo-bleached phenotype was developed four weeks after post-injection. This result indicates that the tobacco rattle virus-induced gene silencing system does work well in MM seedlings. Subsequently, we infiltrated the leaves of *slrnrl1-1* plants with pTRV1/pTRV2 (as a negative control) and pTRV1/pTRV2-*SlRNRL2*. After four weeks, the plants infiltrated with pTRV1/pTRV2 did not show an obvious difference compared to *slrnrl1-1*, while the plants infiltrated with pTRV1/pTRV2-*SlRNRL2* grew very badly, and displayed severe chlorotic leaves, which were not able to recover to green at the late developmental stage (Figure 6F, Supplemental Figure 7B). qRT-PCR analysis revealed that the expression abundance of *SlRNRL2* was indeed down-regulated in plants infiltrated with pTRV1/pTRV2-*SlRNRL2*, compared to *slrnrl1-1* (Figure 6G). All the data demonstrate that SlRNRL1 and SlRNRL2, which possess some redundant functions, are essential for plant growth and development in tomato.

## DISCUSSION

In this study, we systematically studied the biological functions of *SlRNRL1* and *SlRNRL2*, which encode the large subunits of RNR complex of tomato, and demonstrated that the two genes with redundant functions are essential for plant growth and development. Of them, SlRNRL1 plays a more important role than that of SlRNRL2. The loss function of *SlRNRL1* significantly influenced the expression of genes related to DNA metabolic process, cell cycle, photosynthesis, chlorophyll binding, and chloroplast thylakoid membrane (Figure 4C), consequently resulting in young leaf chlorosis and retarded growth, while *SlRNRL2-*knockout mutant *slrnrl2* did not display any visible phenotype compared to wild type.

*SlRNRL1* and *SlRNRL2* of tomato are two genes shared high sequence identity, and their coding proteins were able to interact with SlRNR2 to form RNR complex (Figure 5B). Under the genetic background of *slrnrl1*, *SlRNRL2* expression was markedly upregulated compared to MM (Figure 6E). Corresponding to the increased expression of *SlRNRL2* in the late stage, the chlorosis leaves of *slrnr1* became green. Moreover, knocking down the expression of *SlRNRL2* led *slrnrl1-1* mutant more chlorosis and retarded growth (Figure 6F). All these suggest that *SlRNRL1* and *SlRNRL2* are two homologous genes in the genome of tomato and possess redundant functions. Compared to *slrnrl1*, the loss function of SlRNRL2 did not result a visible phenotype (Figure 6C). This indicates that SlRNRL1 plays a more important role in RNR complex formation than that of SlRNRL2 in tomato. This phenomenon can be explained with their expression difference (high-level expression of *SlRNRL1* and low-level expression of *SlRNRL2*, Figure 5B). *SlRNRL1* and *SlRNRL2* of tomato are very similar to *RNR1* and *RNR3* of yeast. The *RNR1* with a higher expression level is essential for the viability of yeast cells, whereas *RNR3* with lower expression level is not important, and its null mutant does not display any phenotype (Elledge and Davis, 1990). However, overexpression of *RNR3* is able to rescue *RNR1* null mutants. Therefore, we speculate that overexpression of *SlRNRL2* may rescue the phenotype of *slrnrl1*.

Considering of the phenotypes of *slrnrl1* mutants, RNRL1 is directly or indirectly involved in plant growth and development. Similar to Arabidopsis *cls8* (Garton et al., 2007), rice *v3* (Yoo et al., 2009), and Setaria *sistl1* (Tang et al., 2019) mutants, the loss function of SlRNRL1 in tomato results in growth retardation and young leaf chlorosis with decreased chlorophyll contents and chloroplasts number (Figure 1A-1G, 3C-3H). Additionally, the fruit of *slrnrl1* mutants was notably smaller than that of MM, whereas no size difference of siliques was determined between Arabidopsis *cls8-1* and its wild type (Garton et al., 2007). The reason might be as follows. First, the mutation pattern is different. *cls8-1* is a missense substitution mutant changing only one amino acid, while *slrnrl1-1* with 2 bp deletion and *slrnrl1-2* with 1 bp insertion which lead to frame shift and premature termination of SlRNRL1. The mutation of *slrnrl1-1* and *slrnrl1-2* may have a more serious effect on the activities of RNR. Second, the type of fruit is different. Arabidopsis is a silique while tomato is a berry with a lot of flesh.

It has been reported that RNR is crucial to cell cycle progression (Wang and Liu, 2006; Tang et al., 2019). In our study, the ratios of 4C and 8C cells are significantly reduced in *slrnrl1-1* mutant compared to MM (Figure 3B; Supplemental Figure 3C, 3D). We also found that the number of cells in *the slrnrl1-1* mutant was decreased (Figure 3A, Supplemental Figure 3A, 3B). All these suggest that the loss of *SlRNRL1* results in a lower cell division, consequently leading to the smaller leaf and shorter stature of the mutants. The *slrnrl1* mutants exhibited not only a decrease in nuclear genome replication and cell division, but also reduction in the size and number of chloroplasts (Figure 3C-J). We also found photosynthesis was impaired in *slrnrl1-1* (Supplemental Figure 4B, 4C). DEGs enriched in the photosynthesis GO term, and most of them were down-regulated (Figure 4C). As the chloroplast is the factory of plant photosynthesis, the abnormal chloroplasts in *the slrnrl1-1* mutant certainly results in the reduction of photosynthesis. Tomato fruit is a photosynthate sink, its final size relies on the supply of photo-assimilates from leaves. Therefore, the reduced fruit size is a consequent result of the lower cell division rate and impaired leaf photosynthesis in *slrnrl1*.

As shown in Table 1, the expected recombination genotypes except *slrnrl1slrnrl2* have been identified by analyzing more than 300 F2 plants/seeds derived from the cross of *slrnrl1* with *slrnrl2*. This result indicates that the recombination between *SlRNRL1* and *SlRNRL2* occurred in the F2 generation, although the two genes located on the same chromosome with a physical distance of about 46 Mb. The lack of seeds with the *slrnrl1slrnrl2* genotype in the F2 population may be caused by embryo lethality of the double mutant. Deficiency of the large subunit of RNR in the double mutant will block the *de novo* biosynthesis of dNTPs, consequently inhibiting seed formation. Based on that the seeds were able to be developed in the fruits of *slrnrl1* and *slrnrl2*, we speculate that at least one of the two genes (*SlRNRL1* and *SlRNRL2*) is essential for seed development in tomato. Previous studies have demonstrated that the existence of a *de novo* biosynthesis pathway of dNTPs during the embryogenic process and in germinating embryos (Schimpff et al., 1978; Stasolla et al., 2001a, b). Taken together, our data indicate that *SlRNRL1* and *SlRNRL2* are required for the development and growth of tomato.

In conclusion, we demonstrate that the large subunit of RNR complex, encoded by *SlRNRL1* or *SlRNRL2*, is crucial to cell division and chloroplast biogenesis in tomato. Of the two genes, *SlRNRL1* plays a more important role than that of *SlRNRL2* due to its high expression intensity in tomato. Our results also indirectly indicate that the *de novo* biosynthesis of dNTPs is required for seed development of tomato. These findings give new insights into understanding the mechanism of *de novo* biosynthesis of dNTPs in plants.

## Methods

### Plant materials and growth conditions

The *ylc1* (*young leaf chlorosis 1*) mutant was isolated from the EMS-induced mutation population of the tomato cultivar Moneymaker (MM). A wild species, *Sonalum pennellii* (LA0716), was crossed with *ylc1* to construct an F2 population. The *slrnrl1-2* and *slrnrl2* mutants were generated with MM by CRISPR-CAS9 technology and confirmed by sequencing analysis. Seeds were surface sterilized in 70% ethanol for 2 min and 15% commercial bleach for 15 min, and then rinsed with sterilized water three times. The sterilized seeds were germinated on moistened filter paper in a glass petri dish for 7 days. Subsequently, tomato seedlings were grown in growth chambers maintained with 16 h light at 25°C and 8 h dark at 18°C.

### Measurements of physiological characteristics

One-week-old seedlings of MM and *slrnrl1-1/ylc1* grown on moistened filter papers were transferred to the soil for another 2 weeks. The young leaves were collected, and fresh weights were determined. Chlorophyll was extracted in acetone and its content was calculated as described previously (Chen et al., 2018). For the determination of fruit mass, the first three fruits in a plant were harvested and their fresh weight was measured.

Plant height, fresh weight, and dry weight of the mutants and MM were measured after cultivation in Hoagland solution for 3 weeks. Data were collected from at least 3 independent biological replicates.

For the I2–KI dye, the young leaves of MM and *slrnrl1-1* were bleached in a solution with 50% acetone and 50% ethanol overnight, and then dyed in I2–KI solution for 12 h. The samples were then documented by photograph.

### Genetic mapping of the *YLC1* locus

For bulk population sequencing, two DNA pools, LA0716-type pool and *ylc1*-type pool, each with an equal amount of DNA from 50 F2 individuals, derived from *the ylc1* x LA0716 cross, were performed. The two pooled libraries and two parent libraries were prepared and sequenced by Oebiotech Compony (Oebiotech, Shanghai). The △SNP index and the region of the candidate gene were identified as described (Sun et al., 2020).

To exclude irrelevant SNPs from the results of BSA-resequencing, 200 F2 individuals with mutation phenotype were selected and used for the fine mapping of the *YLC1* locus.

### Plasmid construction and plant transformation

For the complementation test, the full length coding sequence (CDS) of *SlRNRL1* was amplified by PCR from tomato cDNA and cloned into the binary vector PBI121. For subcellular localization, the *SlRNRL1-GFP* fusion protein gene was inserted into the vector pJIT163. For the expression pattern of *SlRNRL1*, 1500 bp of genomic DNA upstream of the *SlRNRL1* start codon was amplified and cloned into the pBI121 vector between the *BamH* I and *Hind* III sites. The vector PHSE401 was used for knock-out experiments (Xing et al., 2014). sgRNAs targeting the coding regions of *SlRNRL1* and *SlRNRL2* were designed and cloned into the vector PHSE401 between two *Bsa* I sites.

All constructs generated above were introduced into *A. tumefaciens* strain LBA4404 and then transformed into the tomato genome using *Agrobacterium-*mediated transformation with cotyledon explants (Deng et al., 2018). Transgenic lines were selected based on their resistance to hygromycin B or kanamycin, and confirmed by sequencing and PCR analysis.

### RNA extraction and RT-qPCR

Total RNA was extracted from the tissues using TRIzol Reagent (Invitrogen, USA). After removing genomic DNA by DNase I (Invitrogen, USA), 1 µg total RNA was used to synthesize the first-strand cDNA with M-MLV reverse transcription kit (Invitrogen, USA). RT-qPCR was performed on a LightCycler480 machine (Roche Diagnostics, Switzerland) with SYBR Premix Ex Taq polymerase (TaKaRa, Japan). The relative expression level of target genes was normalized by *Kinase I*. The gene expression abundance is analyzed based on three biological replicates. All primers used for RT-qPCR are listed in Supplemental Table 1.

### Histochemical GUS analysis

The activity of GUS was analyzed as previously described (Wang et al., 2005). Briefly, transgenic lines bearing the *SlRNRL1* promoter/*GUS* fusion construct (p*SlRNRL1*::*GUS*) were incubated at 37℃ overnight with GUS staining solution (100 mM sodium phosphate buffer, pH 7.2, 10 mM EDTA, 0.1% Triton, and 1 mM 5-bromo-4-chloro-3-indolyl-b-D-glucuronic acid). After washing several times using 70% ethanol, the samples were documented by photograph.

### Subcellular localization assay

Arabidopsis protoplasts isolated from leaves of 21-day-old seedlings were transfected with the vector containing *SlRNRL1-GFP* as described previously (Yoo et al., 2007). Briefly, the protoplasts were isolated and harvested from the leaves, digested by cellulase R10 and macerozyme R10, after washing two times using the W5 solution, the protoplasts were re-suspended in MMG solution, and transfected with the corresponding vector through PEG–calcium fusion. After incubation for 12h, the fluorescence signals were detected with a Leica SP8 Confocal microscope system (Leica, Germany).

### Cytology analysis

The cells were counted as described previously (Yuan et al., 2010). Briefly, the chlorophyll of MM and *slrnrl1-1/ylc1* leaves was cleared in the chloral solution (chloral hydrate: glycerol: water, 8:2:1). Then, the unit area was first photographed and the number of palisade cells within the area was counted.

To generate resin-embedded sections, the leaf tissues were fixed in carnoys (acetic acid: ethanol=3:1), then the samples were dehydrated in a gradient ethanol series. Samples were embedded in Technovit® 7100 for sectioning. The Semi-thin sections were stained with 0.25% toluidine blue.

To observe the chloroplast ultrastructure, the 6-day-old first true leaves of MM and *slrnrl1-1* were harvested. Leaf sections were fixed in 2.5% glutaraldehyde at 4°C for 2d and washed with phosphate buffer. The samples were then transferred to 1% OsO_4_. After dehydration in a gradient ethanol series, the samples were embedded in Spurr’s resin prior to ultrathin sectioning. Sections were stained with uranyl acetate and examined with a transmission electron microscope (JET-1400, Japan).

### Flow cytometry

Ploidy level was determined by flow cytometric analysis as described previously with minor modifications (Yuan et al., 2010). The first 6-day-old true leaves were dipped into cold nuclear extraction buffer (20 mM MOPS, pH 5.8, 30 mM sodium citrate, 45 mM MgCl_2_, and 0.1% [v/v] Triton X-100) and chopped with a razor blade. Nuclei were obtained through a 40 µm cell strainer and 500 μl of isolated cells was collected. After staining with 2μg mL^−1^ 4′, 6-diamidino-2-phenylindole (DAPI), the samples were analyzed with a BD FACSCalibur flow cytometer, and 10,000 nuclei were analyzed per experiment.

### The measurement of net photosynthetic rate

The young leaf photosynthesis was measured using the Li-6400 portable photosynthesis system (United States) according to manufacture instructions.

### Protein sequence analysis and phylogeny tree building

The functional domains and motifs were predicted in the Interpro protein prediction database (http://www.ebi.ac.uk/interpro/). Amino acid sequence alignment was done in the MPI Bioinformatics Toolkit (https://toolkit.tuebingen.mpg.de/). The phylogenetic analysis was performed using MEGA4.

### Transcriptome sequencing analysis

The 6-day-old first true leaves of MM and *slrnrl1-1* were harvested for total RNA extraction. cDNA was obtained by reverse transcription through a random N6 primers. Then, the cDNA was sequenced by a GISEQ-500 sequencing instrument. After removing reads containing adapters, reads containing poly-N, and low quality reads from the raw data, high-quality clean reads were obtained. The clean reads were then mapped to the tomato genome using Hisat2 tools (Kim et al., 2015). The gene expression abundances were estimated by fragments per kilo base of the exon model per million mapped fragments (FPKM). For differential expression analysis, clean reads of MM and *slrnrl1-1* were analyzed with the DESeq2 package. The genes with adjusted |log2RPKM*slrnrl1-1*/MM|>1 and *Q-v*alue<0.001 were assigned as differentially expressed genes (DEGs) (Wang et al., 2010). Gene Ontology (GO) enrichment analysis of differentially expressed genes (DEGs) was implemented by the R package (https://en.wikipedia.org/wiki/Hypergenometric-distribution). The qRT-PCR primers for the validation of the RNA-seq results are listed in supplementary Table 1. The genes for heatmap are listed in supplementary Table 2.

### Yeast two-hybrid (Y2H) assay

The Y2HGOLD yeast strain was used Y2H assays. Yeast transformation was conducted with Yeast Transformation System 2 (Clontech NO.630439). All primers used were listed in supplementary Table 1.

### Virus-induced gene silencing (VIGS) in tomato

The VIGS experiment was performed as described previously (Liu et al., 2002). Briefly, *A. tumefaciens* strain GV3101 containing pTRV1 or pTRV2 and its derivatives were grown overnight at 28°C, the *Agrobacterium* cells were harvested, and resuspended in infiltration medium. The bacteria cell density was adjusted to O.D. of 2.0. Equal volumes of *Agrobacterium* strains carrying vectors pTRV1 or pTRV2 and its derivatives were mixed and were injected into the leaves of MM or *slrnrl1-1*.

### Accession Numbers

The sequence data of this article can be found in the SOL Genomics Network Genome (http://solgenomics.net/) under the following accession numbers: *SlRNRL1* (Solyc04g012060), *SlRNRL2* (Solyc04g051350), *SlRNR2* (Solyc01g080210), *Kinase I* (Solyc07g066600).

## Supplemental data

**Supplemental Figure 1.** Cloning of *YLC1*.

**Supplemental Figure 2.** The structure of SlRNRL1 and multiple sequence alignment of the R1 proteins.

**Supplemental Figure 3.** SlRNRL1 is involved in regulating cell cycle progress.

**Supplemental Figure 4.** SlRNRL1 is involved in chloroplast biogenesis and photosynthesis.

**Supplemental Figure 5.** Analysis of the RNA-Seq data of MM and *slrnrl1-1*.

**Supplemental Figure 6.** The Relative expression levels of the genes for validation of RNA-seq result.

**Supplemental Figure 7.** The phenotype of PDS-silenced MM and *slrnrl1-1* plants infiltrated with TRV1/TRV2.

**Supplemental Table 1.** Primers used in this work.

**Supplemental Table 2.** DEGs in mutant phenotype associated GO terms.

## Author contributions

M.-J.G. and H.-L.W. designed and performed the experiments. H.-Q.L. supervised the work. Y.L. and M.C. helped perform the experiments. M.-J.G. H.-Q.L. and H.-L.W. wrote the manuscript.

## Acknowledgements

No conflict of interest declared.

